## Supplemental figures for "Visual and motor signatures of locomotion dynamically shape a population code for feature detection in *Drosophila*"

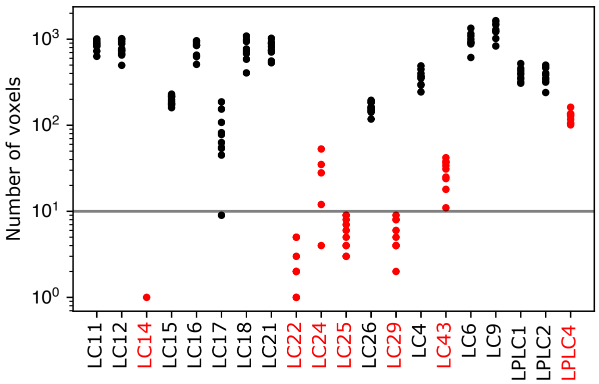


**Figure S1: Number of voxels in each LC/LPLC glomerulus** For each LC/LPLC glomerulus in or near the imaging volume, the number of voxels in each transformed glomerulus mask that lies within the imaging volume is shown for 10 flies. Glomeruli highlighted in red were excluded from this analysis because they were consistently small, or often not included in the volume. The LPLC4 glomerulus was excluded because this glomerulus is co-extensive with the LC22 glomerulus. The horizontal line indicates a threshold we set for further excluding glomeruli from individual flies for being small in that fly's imaging volume. For example, the LC17 glomerulus was sometimes excluded from individual flies because it lies at the bottom of our imaging volume, and brain motion or slight changes in the imaging depth caused the imaged portion of the LC17 glomerulus to be very small.

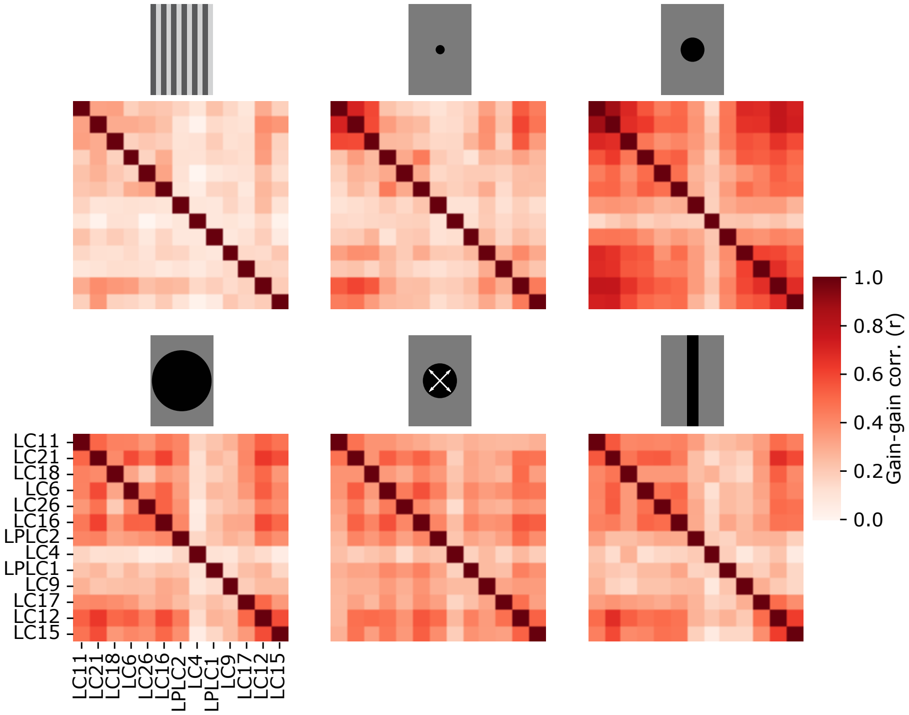


**Figure S2: Trial correlation matrix for each stimulus class** From the trial covariance analysis in Fig. 3, each panel shows the trial correlation matrix for each of the six stimuli shown in the reduced stimulus set for these experiments. Positive correlations are seen for all stimuli, but tend to be higher for stimuli that drive glomerulus responses strongly.

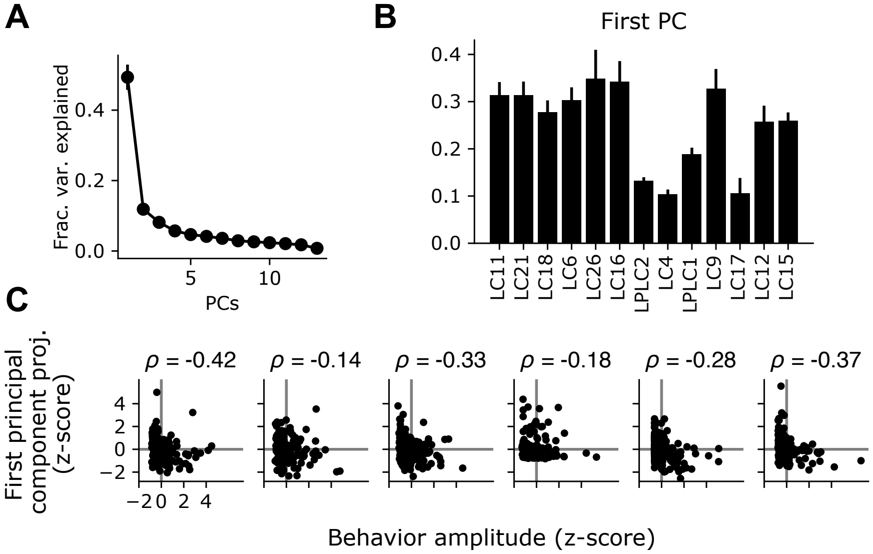


**Figure S3: The shared glomerulus gain factor is negatively correlated with behavior.** (A) Eigenvector decomposition of the covariance matrix reveals a dominant principal component which accounts for a large plurality of the trial-to-trial variance across the population. Points show mean +/- S.E.M. across flies (n=6 flies). (B) The first principal component of the response gain, averaged across flies. (C) For the variance analysis experiments in Fig. 3, we tracked behavior for a subset of the animals tested. For each fly, we z-scored the behavior amplitude across trials as well as the shared gain factor for each trial (equal to the projection of the first PC against the population response matrix). Each plot shows the relationship between the trial-by-trial shared gain factor and the average behavior amplitude for each trial. For every fly, there was a negative correlation between these two measures (mean $\rho$ = -0.23, n=6 flies).

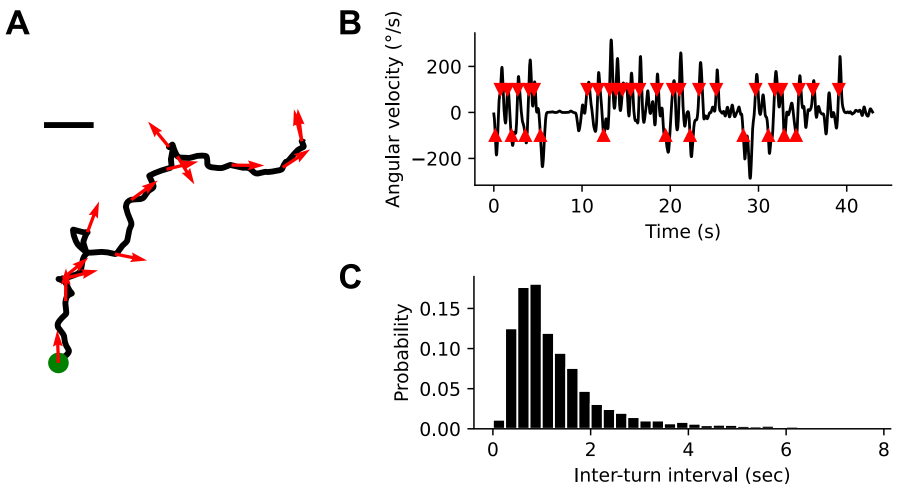


**Figure S4: Natural fly walking consists of punctuated saccadic turns.** (A) Example fly walking trajectory measured in a one square meter arena. Arrows show the fly's heading. Green point indicates start of walking trajectory. Scale bar = 1 cm. (B) Instantaneous angular velocity for the example walking trajectory in (A). Red arrowheads indicate turn events. (C) Histogram of inter-turn intervals shows a peak around 1 second and a long tail.
